# Enhanced neutralization of SARS-CoV-2 XBB sub-lineages and BA.2.86 by a tetravalent COVID-19 vaccine booster

**DOI:** 10.1101/2023.09.14.557682

**Authors:** Xun Wang, Shujun Jiang, Wentai Ma, Xiangnan Li, Kaifeng Wei, Faren Xie, Chaoyue Zhao, Xiaoyu Zhao, Chen Li, Rui Qiao, Yuchen Cui, Yanjia Chen, Jiayan Li, Guonan Cai, Changyi Liu, Zixin Hu, Wenhong Zhang, Mingkun Li, Yanliang Zhang, Pengfei Wang

**Affiliations:** Shanghai Pudong Hospital, Fudan University Pudong Medical Center, Shanghai Institute of Infectious Disease and Biosecurity, State Key Laboratory of Genetic Engineering, MOE Engineering Research Center of Gene Technology, School of Life Sciences, Fudan University, Shanghai, China; Department of Infectious Diseases, Nanjing Hospital of Chinese Medicine Affiliated to Nanjing University of Chinese Medicine, Nanjing, Jiangsu, China; Nanjing Research Center for Infectious Diseases of Integrated Traditional Chinese and Western Medicine, Nanjing, Jiangsu, China; Beijing Institute of Genomics, Chinese Academy of Sciences, and China National Center for Bioinformation, Beijing, China; University of Chinese Academy of Sciences, Beijing, 100049, China; State Key Laboratory of Genetic Engineering, Collaborative Innovation Center for Genetics and Development, School of Life Sciences and Human Phenome Institute, Zhangjiang Fudan International Innovation Center, Fudan University, Shanghai, China; Artificial Intelligence Innovation and Incubation Institute, Fudan University, Shanghai, China; Department of Infectious Diseases, Shanghai Key Laboratory of Infectious Diseases and Biosafety Emergency Response, National Medical Center for Infectious Diseases, Huashan Hospital, Fudan University, Shanghai, China

## Abstract

As the SARS-CoV-2 virus continues to evolve, novel XBB sub-lineages such as XBB.1.5, XBB.1.16, EG.5, HK.3 (FLip), and XBB.2.3, as well as the most recent BA.2.86, have been identified and aroused global concern. Understanding the efficacy of current vaccines and the immune system’s response to these emerging variants is critical for global public health. In this study, we evaluated the neutralization activities of sera from participants who received COVID-19 inactivated vaccines, or a booster vaccination of the recently approved tetravalent protein vaccine in China (SCTV01E), or had contracted a breakthrough infection with BA.5/BF.7/XBB virus. Comparative analysis of their neutralization profiles against a broad panel of 30 SARS-CoV-2 sub-lineage viruses revealed that strains such as BQ.1.1, CH.1.1, and all the XBB sub-lineages exhibited heightened resistance to neutralization than previous variants, however, despite the extra mutations carried by emerging XBB sub-lineages and BA.2.86, they did not demonstrate significantly increased resistance to neutralization compared to XBB.1.5. Encouragingly, the SCTV01E booster vaccination consistently induced robust and considerably higher neutralizing titers against all these variants than breakthrough infection did. Cellular immunity assays also showed that the SCTV01E booster vaccination elicited a higher frequency of virus-specific memory B cells but not IFN-γ secreting T cells. Our findings underline the importance of developing novel multivalent vaccines to more effectively combat future viral variants.

## Introduction

Severe Acute Respiratory Syndrome Coronavirus 2 (SARS-CoV-2) is the virus responsible for causing the disease known as Coronavirus Disease 2019 (COVID-19). With over 769 million confirmed cases and over 6.9 million deaths reported globally as of 6 August 2023^1^, the magnitude of the pandemic remains unprecedented. Despite the World Health Organization’s (WHO) declaration on 5 May 2023^2^, ending the public health emergency of international concern for COVID-19, the journey with the virus is far from over. SARS-CoV-2 remains dynamic, with its genetic code undergoing continual mutations^3^. Consequently, we anticipate the emergence of new variants. The Omicron variant and its subsequent sub-lineages stand out as the most antigenically distinct strains thus far, boasting over 30 mutations within the Spike (S) protein alone^4,5^. Recently, the WHO has identified XBB sub-lineages XBB.1.5, XBB.1.16, and EG.5 as the current Variants of Interest (VOIs)^6^, while several other Omicron sub-lineages, including the most recently identified BA.2.86 with over 30 amino acid changes in S compared with BA.2 and XBB.1.5^7^, are categorized as Variants Under Monitoring (VUMs)^8^. Meticulous tracking of viral variants and timely administration of vaccine boosters for vulnerable groups are still crucial.

The swift development and global deployment of various COVID-19 vaccines have been pivotal in curbing the devastating impact of this severe disease. Several COVID-19 vaccines were approved and rolled out at an unprecedented pace^9^, with a staggering 13.5 billion doses have been administered globally, underlining the monumental scale of this endeavor. The approved vaccines including those utilizing the whole virus as well as those based on specific viral components, such as the protein subunit vaccines and mRNA vaccines^10^. Our previous research^11-14^, along with findings from other investigators^15,16,^ indicated that both primary and booster vaccinations from approved vaccines exhibited reduced efficacy against the Omicron variant and its sub-lineages, particularly over an extended period.

China has initiated the rollout of its first quadrivalent COVID-19 vaccine, SCTV01E developed by Sinocelltech, aiming to tackle a resurging wave of infections driven by multiple SARS-CoV-2 variants. It is an Alpha/Beta/Delta/Omicron variants S-trimer vaccine^17^. Multiple clinical trials, conducted in China and the United Arab Emirates, have assessed its safety, tolerability, immunogenicity, and protective efficacy. Phase 3 clinical data suggest SCTV01E’s effectiveness as a booster shot against Omicron and its subvariants, including XBB, with no significant adverse events reported^18^. The vaccine has been approved for emergency use in China as of March and is now being administered in multiple cities. However, there remains a gap in our understanding of how this multivalent booster vaccine compares in efficacy to other vaccines or to natural immunity acquired through breakthrough infections, particularly in the context of recently emerging viral variants.

## Results

### Neutralization of SARS-CoV-2 main sub-lineages prior to XBB.1.5 by breakthrough infection and SCTV01E booster vaccination sera

In this study, we collected blood sample from individuals at day 14-28 after vaccinated with SCTV01E following three doses of inactivated vaccine (n=21), or from individuals at day 14-28 after infection with BA.5 or BF.7 virus post three doses of inactivated vaccine (n=36), or from those infected with XBB virus post three doses of inactivated vaccine (n=24). We also included 11 individuals who received only with three doses of inactivated vaccine (CoronaVac) as control (Fig. 1A and Table S1). To assess the neutralization potency and breadth of the sera following SCTV01E booster vaccination, we initially assembled a panel of pseudoviruses (PsVs). This panel represented a range of SARS-CoV-2 variants, including the wildtype (WT), B.1.351, B.1.617.2, BA.1, BA.2, BA.5, BF.7, BQ.1.1, CH.1.1, and XBB.1.5. (Fig. 1A). These variants were selected because they had previously circulated at high frequencies.

**Figure. 1:**
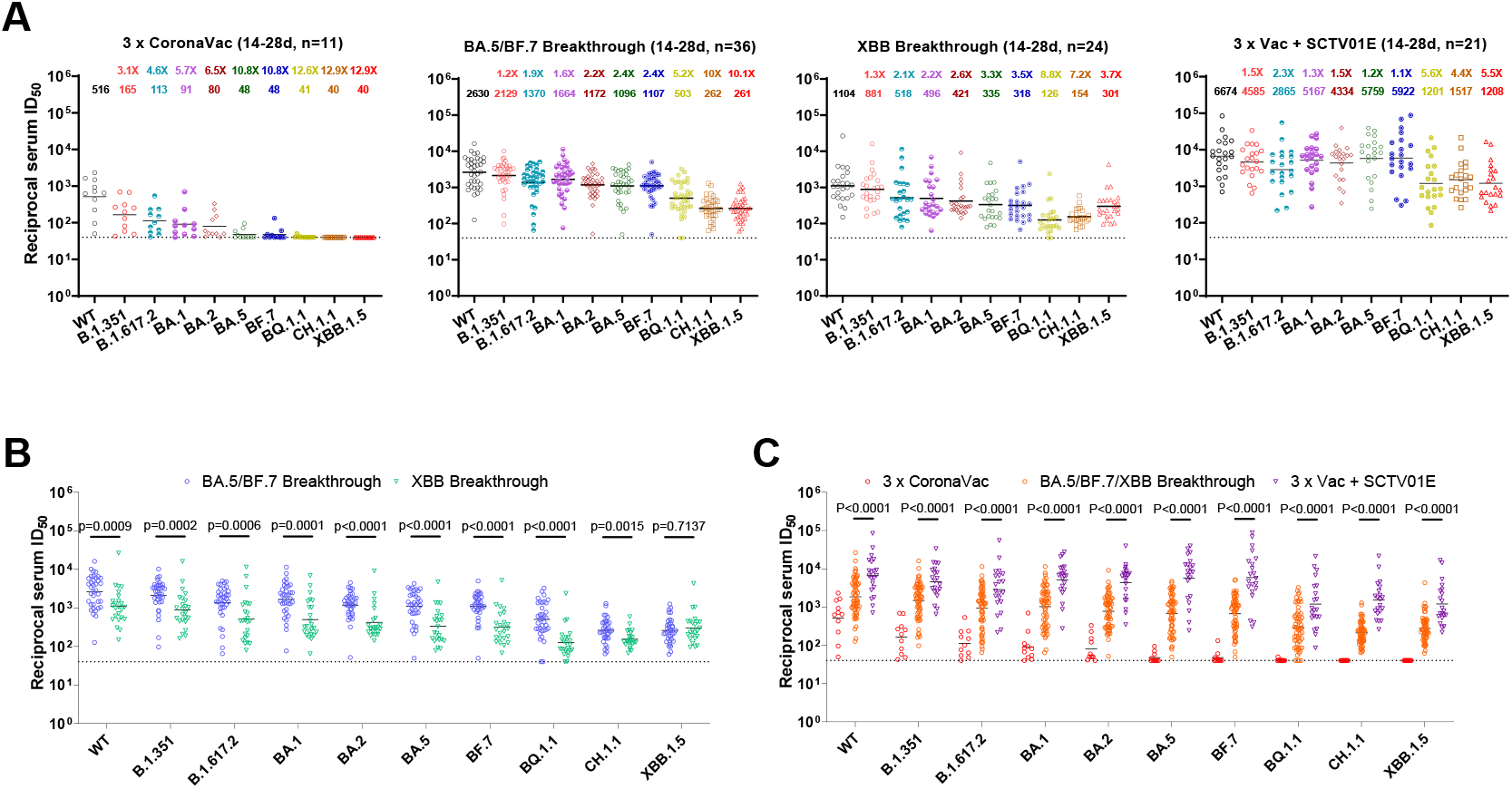
Neutralization of the SARS-CoV-2 main subvariants prior to XBB.1.5 by breakthrough infection and SCTV01E booster vaccination sera. (A) Neutralization of different SARS-CoV-2 variant PsVs by sera collected from individuals at day 14-28 after vaccinated with three doses of CoronaVac, or from individuals at day 14-28 after infection with BA.5/BF.7 virus post three doses of inactivated vaccine, or from those infected with XBB virus post three doses of inactivated vaccine, or from individuals at day 14-28 after vaccinated with SCTV01E following three doses of CoronaVac. (B) In parallel comparison of neutralization GMTs against distinct Omicron subvariants by sera collected from individuals with different SARS-CoV-2 variants breakthrough infections. (C) In parallel comparison of neutralization GMTs against distinct Omicron subvariants by sera collected from individuals with BA.5/BF.7/XBB breakthrough infections, or with SCTV01E booster vaccinations. For all panels, values above the symbols denote geometric mean titer and the fold-change was calculated by comparing the titer to WT. P values were determined by using Multiple Mann-Whitney tests. The horizontal dotted line represents the limit of detection of 40. WT: wild-type.

Consistent with our previous findings^14^, individuals who received a three-dose vaccination series demonstrated detectable geometric mean titers (GMTs) of neutralizing antibodies against the WT virus, but the GMTs for B.1.351, B.1.617.2, BA.1 and BA.2 showed significant reductions compared to WT, while the neutralizing titers against the BA.5, BF.7, BQ.1.1, CH.1.1, and XBB.1.5 were reduced to nearly undetectable levels. In individuals who experienced breakthrough infections with the BA.5 or BF.7 variants, high levels of neutralizing titers were observed against variants that preceded BA.5 and BF.7, with GMTs consistently exceeding 1000. However, neutralizing titers against the BQ.1.1, CH.1.1, and XBB.1.5 variants were substantially diminished (Fig. 1A). Compared to those with BA.5/BF.7 breakthrough infections, individuals who had XBB breakthrough infections showed similar neutralizing titers against the XBB.1.5 variant; however, their neutralizing titers against other variants were significantly lower (Figs. 1A and 1B). This could suggest that the XBB variants possess lower antigenicity than BA.5/BF.7. It is noteworthy that individuals who received the SCTV01E booster vaccination exhibited the highest neutralizing activity against all tested SARS-CoV-2 variants. In the SCTV01E booster group, the GMT for neutralization against the WT virus was 6674. The GMTs against other variants, such as B.1.351, B.1.617.2, BA.1, BA.2, BA.5, and BF.7, ranged from 4585 to 5922, representing only a modest 1.1-to 2.3-fold reduction compared to the WT. Even for the BQ.1.1, CH.1.1, and XBB.1.5 variants, which demonstrated greater resistance to neutralization elicited by the booster (a 4.4-to 5.6-fold reduction), the neutralization titers against these variants remained above 1000 (Fig. 1A). When compared to both the three-dose CoronaVac group and the combined breakthrough infection group (comprising individuals with BA.5/BF.7 and XBB breakthrough infections), the SCTV01E booster group exhibited significantly higher neutralization titers against all the tested SARS-CoV-2 variants (all p-values < 0.0001) (Fig. 1C).

### Neutralization of emerging XBB sub-lineages and BA.2.86 by breakthrough infection and SCTV01E booster vaccination sera

The above results suggested that sera from individuals who received the SCTV01E booster vaccination maintained effective neutralization capabilities against a range of SARS-CoV-2 variants, extending up to the XBB.1.5 variant. However, the viral landscape is evolving, with several new XBB sub-lineages appearing, each carrying additional mutations (Fig. 2A). As of May 2023, XBB.1.5 continued to be the dominant SARS-CoV-2 variant, but there was a noticeable trend of newly emerging variants that began to challenge XBB.1.5’s predominance (Figs. 2B). Notably, the XBB.1.16 subvariant has been gaining momentum worldwide, particularly in countries such as India, the United Kingdom and the United States. Another emerging XBB subvariant, EG.5.1, is also spreading swiftly across the globe, predominantly in countries like China, France, and Singapore (Figs. 2B and S1). This subvariant evolved from Omicron XBB.1.9.1 (with same S sequence as XBB.1.5) and features an additional F456L substitution in the Receptor Binding Domain (RBD) of the S protein, as well as a Q52H mutation in the N-Terminal Domain (NTD) of the S. Yet another emergent subvariant, HK.3, includes a novel L455F substitution in addition to EG.5.1’s F456L, a combination referred to as the “FLip” mutation (Fig. 2C). Sub-lineage XBB.2.3, likely originating in India, has also expanded to multiple countries, including the United States and Singapore (Figs. 2B and S1). Apart from these XBB sub-lineages, BA.2.86, a subvariant that evolved from the once-dominant Omicron BA.2 lineage — now virtually extinct — has garnered significant attention. This is not only due to its unique evolutionary trajectory but also because its genetic sequence includes over 30 amino acid changes in the spike protein, when compared with BA.2 and XBB.1.5 (Figs. 2A and 2D). To further test the protective efficacy of the SCTV01E vaccine against these newly emerging SARS-CoV-2 variants, we constructed a panel of PsVs representing XBB.1.5, XBB.1.16, EG.5, XBB.2.3 and their selected descendent sub-lineages that either have a relatively high prevalence in the population (accounting for more than 1% of the sequences in the past four months) or newly emerged with a notable growth advantage over others (relative growth advantage >= 10%, https://cov-spectrum.org/), as well as BA.2.86 (Fig. 2B-D), for serum neutralization test. A thorough database search revealed an additional 174 lineages that shared identical S protein sequences with our selected lineages. Consequently, this panel of viruses we included here encompasses approximately 80% of the total sequence shared in the past four months (Fig. 2B, from May to August: 80.8%, 78.5%, 75.8%, 74.8%).

**Figure. 2:**
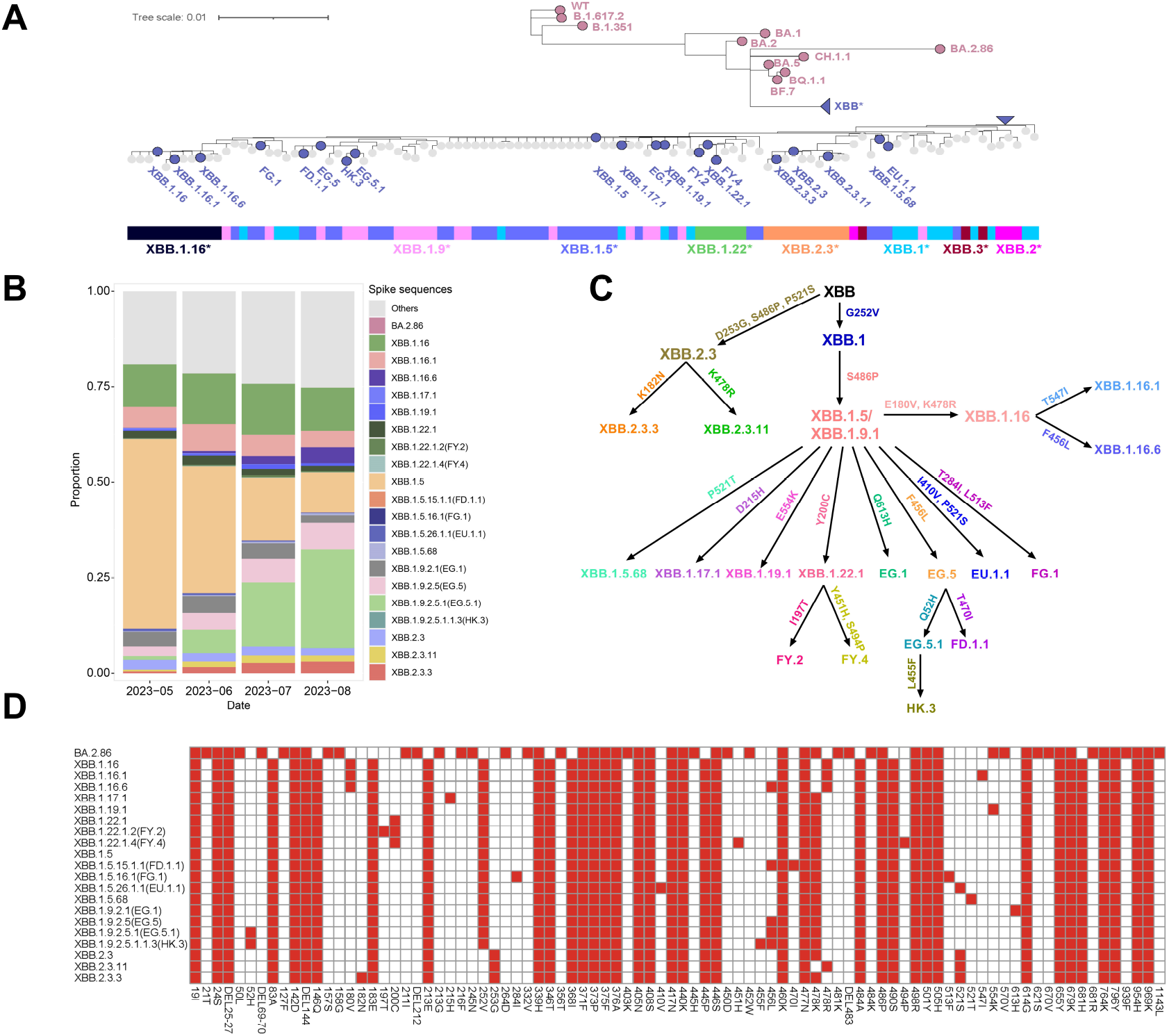
Characteristics of the recently emerging SARS-CoV-2 XBB subvariants. (A) Phylogenetic tree of the SARS-CoV-2 variants and XBB subvariants. The phylogenetic tree of the 21 selected lineages and other XBB sub-lineages with a minimum sequence number of 100. The consensus S mutations of each lineage were obtained from the outbreak.info database with a threshold of 75%. The amino acid sequence of the S protein of each lineage was obtained by replacing each mutation on the reference sequence (EPI_ISL_402124), lineages that shared the identical sequence were merged. Phylogenetic analysis was performed with the MEGA X (v10.1) using the minimum evolution method. (B) The dynamics of the proportion of the 21 selected lineages and other lineages that share the same S sequence. The proportion of other lineages that share the same S sequence is included within each selected lineage. Data was obtained from the RCoV19 database on 31 August 2023. (C) Schematic relationship between XBB-lineage variants in this study. Arrows denote direct relationships between variants with the corresponding S mutations written along them. (D) The S mutations possessed by the additional selected lineages. Mutation data was obtained from the outbreak.info database with a threshold of 75%.

In a similar manner to the methods described earlier, we included serum samples from the BA.5/BF.7 breakthrough infection and the XBB breakthrough infection groups for comparative analysis. As shown in Figs. 3A-3C, for the two breakthrough infection groups and the SCTV01E booster vaccination group, they all exhibited very similar degree of neutralizing titers against the distinct emerging XBB sub-lineages. When compared to the ancestor subvariant XBB.1.5/XBB.1.9.1, the recently surging subvariants were only modestly (<2-fold) more resistant to neutralization by the serum samples. Interestingly, despite its high mutational load, the BA.2.86 variant did not demonstrate greater resistance to neutralization than did XBB.1.5. We then conducted a parallel comparison of neutralizing activities against all these XBB sub-lineages between the BA.5/BF.7 and XBB breakthrough infection groups. The latter group exhibited marginally higher neutralization titers, although the difference was not statistically significant, with the exception of the EG.5.1 subvariant (Fig. 3D). Consequently, we aggregated the data from both breakthrough infection groups and compared their GMTs against those of the SCTV01E booster vaccination group. Once again, the SCTV01E group displayed significantly higher neutralization titers against all tested SARS-CoV-2 variants, with all p-values falling below 0.0001 (Fig. 3E). Taken together, despite the additional mutations carried by emerging XBB sub-lineages and BA.2.86, they did not manifest substantially enhanced resistance to neutralization. Encouragingly, the SCTV01E booster vaccination consistently induced robust neutralizing titers against all these variants, underscoring its potential effectiveness against a broad spectrum of SARS-CoV-2 subtypes.

**Figure. 3:**
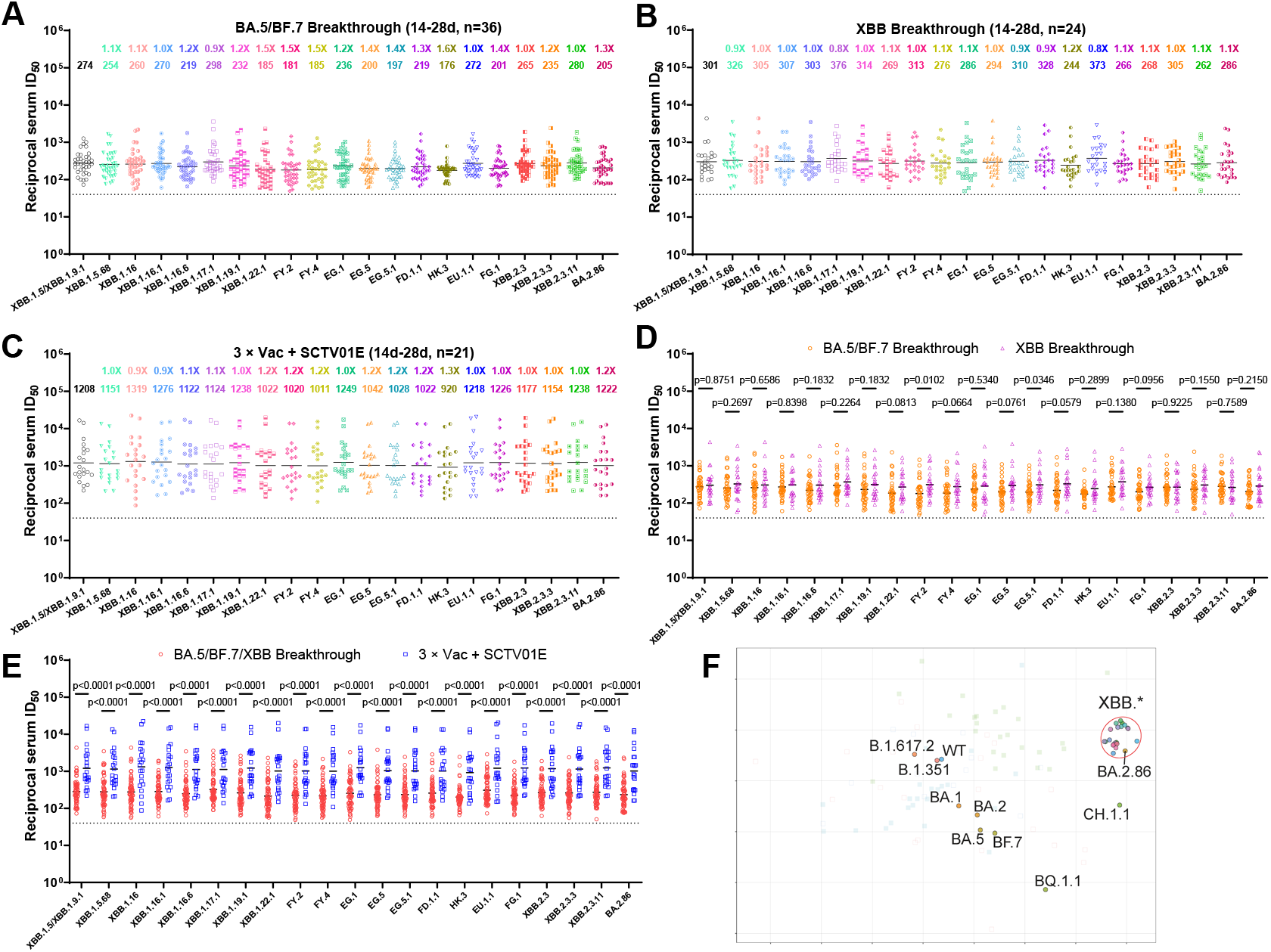
Neutralization of emerging XBB sub-lineages and BA.2.86 by breakthrough infection and SCTV01E booster vaccination sera. Neutralization of different SARS-CoV-2 variant PsVs by sera collected from individuals from individuals at day 14-28 after infection with BA.5 or BF.7 virus post three doses of inactivated vaccine (A), or from those infected with XBB virus post three doses of inactivated vaccine (B), or from individuals at day 14-28 after vaccinated with SCTV01E following three doses of inactivated vaccine (C). (D) In parallel comparison of neutralization GMTs against distinct XBB subvariants by sera collected from individuals with different SARS-CoV-2 variants breakthrough infections. (E) In parallel comparison of neutralization GMTs against distinct XBB subvariants by sera collected from individuals with BA.5/BF.7/XBB breakthrough infections, or with SCTV01E booster vaccinations. (F) Antigenic map based on all the breakthrough infection and SCTV01E booster serum neutralization data from Figs. 1 and 3. Virus positions are represented by closed circles whereas serum positions are shown as open or closed squares. Both axes represent antigenic distance with one antigenic distance unit (AU) in any direction corresponding to a 2-fold change in neutralization ID_50_ titer.

We extended our analysis to measure binding antibody titers, focusing on anti-RBD and anti-S IgG antibodies against the WT, BA.5, and XBB.1.16 viruses. Notably, the SCTV01E group exhibited markedly higher RBD and S binding antibody responses when compared to the BA.5 breakthrough group (Fig. S2A). Further, we explored the correlation between binding and neutralization antibody titers. A consistent correlation was observed when comparing binding antibodies and neutralization titers against both WT and BA.5 viruses. However, no such correlation was evident for the XBB.1.16 subvariant (Fig. S3B). These findings suggest that the interplay between binding and neutralizing antibody responses may become less predictable as the virus evolves, particularly against highly mutated variants. To visualize and quantify the antigenic distances among the various SARS-CoV-2 variants, we utilized our comprehensive set of breakthrough infection and SCTV01E booster neutralization data to construct an antigenic map, as illustrated in Fig. 3F. Interestingly, the SARS-CoV-2 WT virus and its early variants such as Beta (B.1.351) and Delta (B.1.617.2) appear antigenically similar, forming a close-knit cluster. Early Omicron subvariants, including BA.1, BA.2, BA.5, and BF.7, segregate into another distinct cluster. Meanwhile, more recent variants like BQ.1.1, CH.1.1, and the XBB sub-lineages demonstrate considerable antigenic drift, distancing themselves from both the WT and early Omicron clusters. Intriguingly, our data reveal that BA.2.86 aligns closely with the XBB sub-lineages, reinforcing our earlier findings.

### B- and T-Cell responses following breakthrough infection and SCTV01E booster vaccination

After assessing the antibody responses elicited by the SCTV01E booster vaccine, we turned our attention to comparing B and T cell responses among those who received the SCTV01E booster, a full course of the CoronaVac vaccine, and those who had breakthrough infections with the BA.5 variant. Flow cytometry was employed to quantify peripheral blood SARS-CoV-2-specific B and T cells in these groups (Fig. S3). For the B cell responses against the three targeted viruses (WT, BA.5, and XBB.1.16), we observed a significant increase in the frequency of RBD-specific memory B cells (MBCs) in individuals who had a breakthrough infection with the BA.5 variant. Importantly, this frequency was even higher in those who received the SCTV01E booster, as compared to those who had only completed a three-dose regimen of the CoronaVac vaccine, in consistence with our serological findings. These results suggest that the tetravalent protein-based SCTV01E booster vaccine significantly enhances memory B cell immunity against a range of SARS-CoV-2 variants (Fig. 4A).

**Figure. 4:**
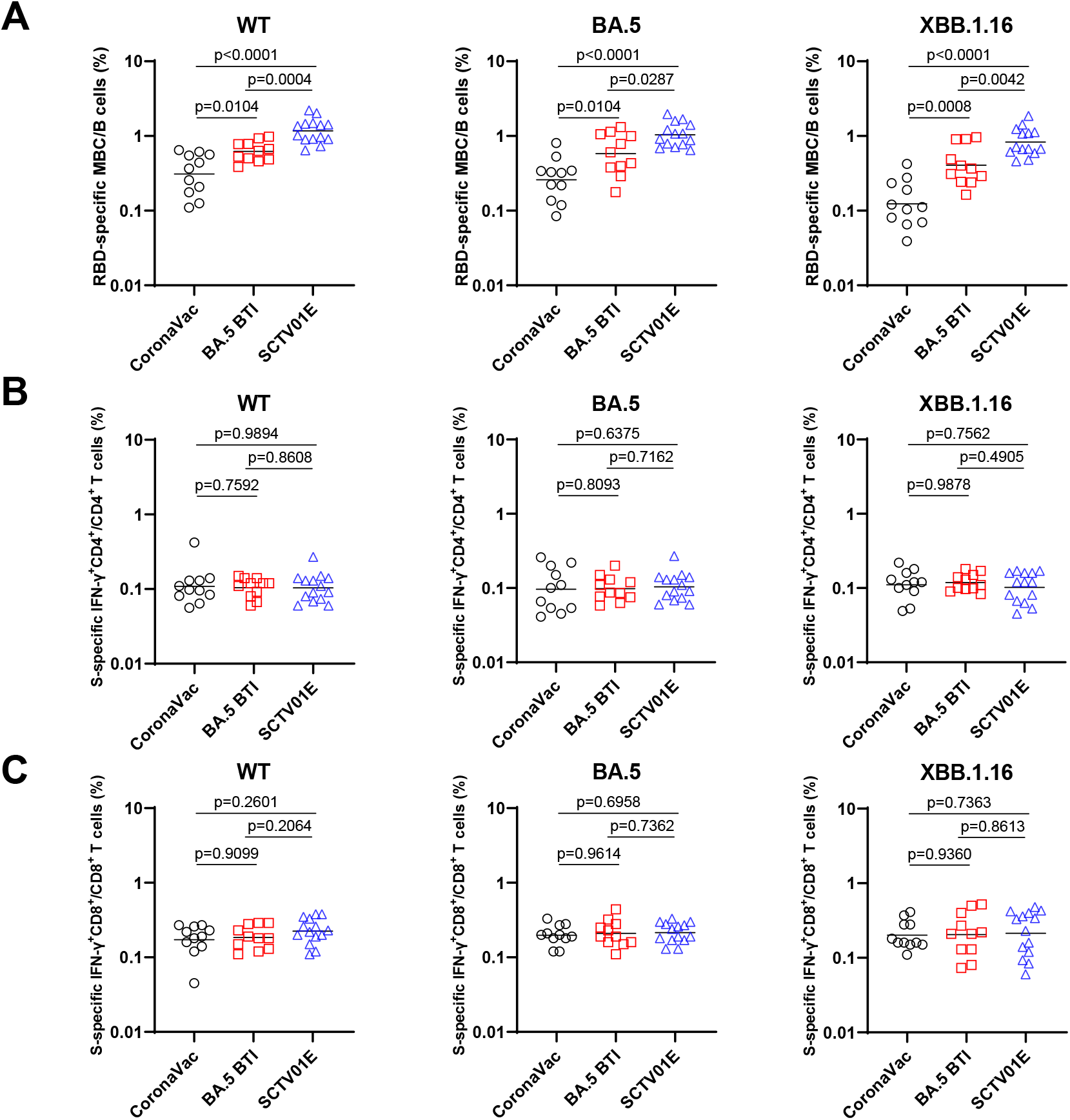
B- and T-Cell responses following breakthrough infection and SCTV01E booster vaccination. (A) The frequencies of WT/BA.5/XBB.1.16 RBD-specific MBCs from individuals with BA.5 breakthrough infections, or with SCTV01E booster vaccinations. Comparison of WT/BA.5/XBB.1.16 S-specific IFN-γ+CD4+T cells (B) or IFN-γ+CD8+T cells (C) between BA.5 breakthrough infections and SCTV01E booster vaccinations groups. P values were calculated using Wilcoxon signed-rank test for group comparisons. RBD, receptor-binding domain; MBCs, memory B cells; IFN-γ, interferon-gamma.

Understanding the role of SARS-CoV-2-specific T-cell responses in convalescent individuals and vaccine recipients is crucial for controlling viral infections and preventing severe symptoms^19^. However, the potential of the SCTV01E booster vaccine to enhance T-cell responses remains undetermined. To explore this, we utilized SARS-CoV-2 WT, BA.5, and XBB.1.16 S trimers to measure differences in S-specific IFN-γ secretion across the same three groups. For T-cell responses, we found no significant differences in the proportions of S-specific IFN-γ-producing CD4+ T cells across the three groups (Fig. 4B). Similarly, the frequency of IFN-γ-producing CD8+ T cells did not significantly differ among these groups (Fig. 4C).

In conclusion, our data suggest that the SCTV01E booster vaccine does not confer a marked advantage in eliciting stronger CD4+ and CD8+ T-cell responses compared to either a three-dose regimen of inactivated vaccine or breakthrough infection with BA.5. Therefore, while the SCTV01E booster significantly enhances antibody and B-cell responses, its impact on T-cell immunity appears to be less distinct.

## Discussion

More than three and a half years into the COVID-19 pandemic, the ongoing evolution and proliferation of new SARS-CoV-2 variants are likely to pose significant global public health challenges for years to come^20^. The virus’s continuous mutations and recombinations have given rise to the XBB variant, along with numerous sub-lineages, necessitating ongoing surveillance and assessments^21^. At present, the global landscape of the virus comprises a diverse set of XBB subvariants, including XBB.1.5, XBB.1.16, EG.5, XBB.2.3, and their subsequent derivatives^22^. Traditional prototype vaccines and variant-specific monovalent vaccines have been shown to have limited cross-neutralizing capabilities against these evolving strains, as evidenced by previous studies^23-27^. Given that several new COVID-19 vaccines have received emergency use authorization, evaluating their immunological effectiveness is crucial. In this study, we recruited individuals who received the S protein-based tetravalent vaccine SCTV01E as a booster shot and also included participants with diverse vaccination and infection histories for comparison. To assess their serum-neutralizing activities, we assembled what is, to the best of our knowledge, the most comprehensive panel of SARS-CoV-2 variants. This panel has a total of 30 variants including both that were widely disseminated in the past and the currently circulating XBB sub-lineages, as well as the most recently identified BA.2.86.

While early variants like B.1.351 and B.1.617.2 exhibited moderate resistance to neutralization and early Omicron subvariants such as BA.1, BA.2, BA.5, and BF.7 showed increased resistance, the most significant resistance has been observed in BQ.1.1, CH.1.1, and all XBB sub-lineages. This substantial resistance could be attributed to the marked antigenic divergence between the BQ/CH/XBB and previous strains. Interestingly, when comparing among all the XBB subvariants and the recently identified BA.2.86, we found that they did not display significantly enhanced resistance to neutralization relative to XBB.1.5. Even for recently emerging and concerning subvariants like EG.5.1 and HK.3 (FLip), they did not exhibit notably increased resistance to neutralization compared to their parent strain, XBB.1.5. In fact, according to the antigenic map we constructed, all XBB subvariants and BA.2.86 clustered relatively closely together, although they were far removed from other viruses. This suggests that despite their numerous mutations, these strains have not developed dramatically enhanced capabilities to evade neutralization.

In comparison to the BA.5/BF.7 breakthrough infection group, the XBB breakthrough infection group displayed lower cross-reactivity against all major pre-XBB strains. However, their neutralizing titers against the various XBB sub-lineages were fairly similar across both breakthrough infection groups. While breakthrough infections elicited much higher neutralizing antibody responses than a three-dose regimen of an inactivated vaccine, the SCTV01E booster vaccination yielded the highest neutralization activities across all cohorts for all tested viruses. This suggests that individuals who have been fully vaccinated and then boosted with this novel vaccine are likely to have enhanced protection against emerging XBB subvariants as well as BA.2.86. Although further studies are required to determine the duration of this protective efficacy, our research indicates that multivalent vaccines can broaden the diversity of antibody responses, thereby improving cross-strain protection.

In addition, we conducted a preliminary investigation into cellular immune responses using flow cytometry to assess the frequency of RBD-specific MBCs and S-specific IFN-γ secreting T cells. In line with our serological findings on neutralizing antibodies, both SCTV01E booster vaccination and breakthrough infection substantially enhanced MBC immunity against a range of SARS-CoV-2 variants. This heightened MBC response may be instrumental in generating robust levels of neutralizing antibodies capable of thwarting reinfections by newly emergent variants. These results lend credence to the idea that repeated exposure to viral antigens, whether through natural infection or booster immunization with a novel vaccine, can fortify our B-cell immune response against evolving strains^28-30^. Moreover, a robust SARS-CoV-2-specific T-cell response, whether induced by vaccination or natural infection, can effectively eradicate infected cells, curb further viral spread, and guard against severe disease^31^. However, our data did not reveal any statistically significant differences in viral-specific T-cell responses across the cohorts, regardless of whether they had received booster vaccinations or had a history of hybrid immunity. It’s worth noting that our analysis of B- and T-cell responses is preliminary. Future studies should delve deeper, exploring factors like the dynamic profiles of plasma cells, subtypes of MBCs, follicular helper T cells, and regulatory T cells.

In summary, although the SARS-CoV-2 virus continues to evolve and enhance its ability to evade immunity, the currently emerging XBB sub-lineages and BA.2.86 have not drastically diverged antigenically from the original XBB cluster. The markedly higher neutralizing titers induced by SCTV01E vaccination, compared to prototype vaccines and natural infection, underscore the importance of developing novel multivalent vaccines to more effectively combat future viral variants.

## DATA AVAILABILITY

All the data are provided in the main or the supplementary figures.

## ACKNOWLEDGMENTS

This study was supported by funding from the National Natural Science Foundation of China (32270142 to P.W.), Shanghai Rising-Star Program (22QA1408800 to P.W.), the Program of Science and Technology Cooperation with Hong Kong, Macao and Taiwan (23410760500 to P.W.), Nanjing Research Center for Infectious Diseases of Integrated Traditional Chinese and Western Medicine (YBZX2022 to Y.Z.), and NATCM’s Project of High-level Construction of Key TCM Disciplines. Pengfei Wang acknowledges support from Open Research Fund of State Key Laboratory of Genetic Engineering, Fudan University (No. SKLGE-2304) and Xiaomi Young Talents Program.

## AUTHOR CONTRIBUTIONS

P.W., Y.Z., and M.L. designed and supervised the study; X.W., S.J., and W.M. performed the experiments with help from X.L., C.Z., X.Z., C.L., R.Q., Y. Cui, Y. Chen., J.L., G.C., C.L., and Z.H.; S.J., K.W., F.X., W.Z., and Y.Z. provided critical materials; P.W., Y.Z., M.L., X.W., S.J., and W.M. analyzed the data and wrote the manuscript. All authors reviewed, commented, and approved the manuscript.

## DECLARATION OF INTERESTS

All authors declare no conflict of interest.

## Materials and methods

### Serum samples

Sera from individuals who received three doses of inactivated vaccine (CoronaVac) or boosted with tetravalent protein-based vaccinee (SCTV01E) after receiving three doses of inactivated vaccine were collected at Nanjing Hospital of Chinese Medicine 14-28 days after the final dose. 22 individuals who were infected with BA.5 variant, 14 individuals who were infected with BF.7 variant, and 24 individuals who were infected with XBB variant after receiving three doses of inactivated vaccine were recruited at the Nanjing Hospital of Chinese Medicine. For all COVID-19 participants, the clinical diagnosis criteria were based on the ninth National COVID-19 guidelines. The SARS-CoV-2 infection of all the subject was confirmed by Polymerase Chain Reaction (PCR) and sequencing. All participants involved in this study had mild symptoms. Their baseline characteristics are summarized in Table S1. All the participants provided written informed consents. All collections were conducted according to the guidelines of the Declaration of Helsinki and approved by the ethical committee of Nanjing Hospital of Chinese Medicine Affiliated to Nanjing University of Chinese Medicine (number KY2021162).

### Cell lines

Expi293F cells (Thermo Fisher Cat# A14527) were cultured in the serum free SMM 293-TI medium (Sino Biological Inc.) at 37 °C with 8% CO_2_ on an orbital shaker platform. HEK293T cells (Cat# CRL-3216), Vero E6 cells (cat# CRL-1586) were from ATCC and cultured in 10% Fetal Bovine Serum (FBS, GIBCO cat# 16140071) supplemented Dulbecco’s Modified Eagle Medium (DMEM, ATCC cat# 30-2002) at 37°C, 5% CO_2_. I1 mouse hybridoma cells (ATCC, cat# CRL-2700) were cultured in Eagle’s Minimum Essential Medium (EMEM, ATCC cat# 30-2003)) with 20% FBS.

### Construction and production of variant pseudoviruses

Plasmids encoding the WT (D614G) SARS-CoV-2 spike and Omicron sub-lineage spikes were constructed. HEK293T cells were transfection with the indicated spike gene using Polyethylenimine (Polyscience). Cells were cultured overnight at 37°C with 5% CO_2_ and VSV-G pseudo-typed ΔG-luciferase (G*ΔG-luciferase, Kerafast) was used to infect the cells in DMEM at a multiplicity of infection of 5 for 4 h before washing the cells with 1×DPBS three times. The next day, the transfection supernatant was collected and clarified by centrifugation at 3000g for 10 min. Each viral stock was then incubated with 20% I1 hybridoma (anti-VSV-G; ATCC, CRL-2700) supernatant for 1 h at 37 °C to neutralize the contaminating VSV-G pseudotyped ΔG-luciferase virus before measuring titers and making aliquots to be stored at −80 °C.

### Pseudovirus neutralization assays

Neutralization assays were performed by incubating pseudoviruses with serial dilutions of sera, and scored by the reduction in luciferase gene expression. In brief, Vero E6 cells were seeded in a 96-well plate at a concentration of 2×10^4^ cells per well. Pseudoviruses were incubated the next day with serial dilutions of the test samples in triplicate for 30 min at 37 °C. The mixture was added to cultured cells and incubated for an additional 24 h. The luminescence was measured by Luciferase Assay System (Beyotime). IC_50_ was defined as the dilution at which the relative light units were reduced by 50% compared with the virus control wells (virus + cells) after subtraction of the background in the control groups with cells only. The IC_50_ values were calculated using nonlinear regression in GraphPad Prism.

### Antigenic Cartography

The constructed antigenic map was based on serum neutralization data utilizing the antigenic cartography methods, which are implemented in the Racmacs package (https://acorg.github.io/Racmacs/). The antigenic map was generated in R with 10000 optimization steps and other default parameters in a 2-dimesional space. The distances between positions of sub-lineages and serum on the antigenic map were optimized so that distances approach the fold decreases in neutralizing ID_50_ titer, relative to the maximum titer for each serum. Each unit of distance in arbitrary directions in the antigenic map represents a 2-fold change in the ID_50_ titer.

### Protein expression and purification

SARS-CoV-2 S2P S WT/BA.5/XBB.1.16 trimer proteins and WT/BA.5/XBB.1.16 RBD (aa319-541) were separately cloned into mammalian expression vector pCMV3 with an 8 × His tag and an Avi-tag at the C terminus. Proteins were expressed in Expi293F cells by transfection of S trimer or RBD expressing plasmids using Polyethylenimine and culture at 37 °C with shaking at 125 rpm and 8% CO2. On day 5, the supernatant was collected by centrifugation at 4000 × g for 10 min, then incubated with Ni-NTA beads (Invitrogen) for 1 h at room temperature. The beads were washed with ten column volumes of 20 mM imidazole in PBS, then eluted with 500 mM imidazole in PBS. The purified protein was buffer exchanged into plain PBS and concentrated by 10 kDa cutoff column and purity was assessed by OD280 and SDS-PAGE. For enzymatic biotinylation, the Avi-tagged RBD at 1 mg/mL was incubated with 10 mM ATP, 10 mM magnesium acetate, 50 μM D-biotin, 50mM Bicine, 19.2 μg/mL in house-purified BirA in a 1 mL reaction mix and incubated at 30 °C for 5 h.

### Memory B cells detection by flow cytometry

PBMCs were isolated by PBMC isolation kit (Solarbio) according to the manufacturer’s instructions. For SARS-CoV-2-specific B cell detection, approximately 1.0×10^6^ PBMCs were firstly incubated with LIVE/Dead Zombie Aqua Dye (77143, BioLegend; 1 μL/well) for 15min at 4°C. For RBD staining, biotinylated SARS-CoV-2 Spike RBD protein was mixed with Streptavidin-BV421 (405225, BioLegend) at a 4:1 molar ratio for 1 hour at 4 °C to obtain the antigen probe. After being washed and resuspended with Fluorescence-Activated Cell Sorting (FACS) buffer (PBS + 2% FBS), PBMCs were prepared into a cell suspension, added to plates (100 μL/well), and then stained for 30 minutes at 4 °C using antigen probe (1.5 μL/well) and the following conjugated antibodies: APC anti-human CD3 (300412, BioLegend; 1 μL/well), PerCP/Cyanine 5.5 anti-human CD19 (302230, BioLegend; 1 μL/well), PE anti-human CD27 (302808, BioLegend; 1 μL/well). After being stained, the cells were washed and resuspended in 200 μL FACS buffer. The full gating strategy is illustrated in Supplementary Figure 3A.

### Detection of T cells by flow cytometry

For SARS-CoV-2-specific T cell detection, after being washed and resuspended with the complete medium (RPMI-1640 (Giboco, 11875093) + 10% FBS), approximately 2.0×10^6^ PBMCs were prepared into a cell suspension, added to plates (250 μL/well), and then incubated with CD28 Monoclonal Antibody (CD28.2) (302934, BioLegend; 0.5 μL/well), CD49d (Integrin alpha 4) Monoclonal Antibody (9F10), (304340, BioLegend; 0.25 μL/well), Monensin (420701, BioLegend; 0.25 μL/well) and 500 μg/mL SARS-CoV-2 spike trimer (2 μL/well) overnight at 37 °C with 5% CO2. After being washed and resuspended with FACS buffer, cells were prepared into a cell suspension, added to plates (50 μL/well), and then stained for 30 minutes at 4 °C using the following antibodies: Brilliant Violet 711™ anti-human CD4 (344648, BioLegend; 1 μL/well), APC anti-human CD3 (300412, BioLegend; 1 μL/well), and PE/Dazzle™ 594 anti-human CD8 (301058, BioLegend; 1 μL/well). After washing with FACS buffer, the cells were stained for 20 minutes at 4 °C using a Cyto-Fast™ Fix/Perm Buffer Set (426803, BioLegend; 100 μL/tube). After being washed and resuspended with Cyto-Fast™ perm wash solution, cells were prepared into a cell suspension, added to plates (50 μL/well), and then stained for 20 minutes at 4 °C using the following antibodies: Brilliant Violet 605™ anti-human interferon-gamma (IFN-γ) (506542, Biolegend; 1 μL/well). After being stained, the cells were washed and resuspended in 200 μL FACS buffer. FlowJo software was used for the analysis of the B and T cell populations. The full T cells gating strategy is illustrated in Supplementary Figure 3B.

### ELISA

We coated 100 ng per well of S trimer or 100 ng per well of RBD onto ELISA plates at 4 °C overnight. The ELISA plates were then blocked with 300 μl blocking buffer (3% BSA in PBS) at 37 °C for 2 h. Afterwards, sera were serially diluted using dilution buffer (3% BSA in PBS), incubated at 37 °C for 1 h. Next, 100 μl of 2000-fold diluted goat anti-human IgG (H+L) antibody (Beyotime) was added into each well and incubated for 1 h at 37 °C. The plates were washed between each step with PBST (0.5% Tween-20 in PBS). Finally, the TMB substrate (Beyotime) was added and incubated before the reaction was stopped using 1 M sulfuric acid. Absorbance was measured at 450 nm. OD450 readings were normalized by subtracting the average of negative control wells and finally dividing by the average maximum signal for each unique coating protein in each experiment. Normalized OD450 data from biological replicates was combined and fit with a three-parameter logistic model.

### Prevalence of selected lineages

Sequence metadata was obtained from the RCoV19 database^32^ on 31 August 2023. Lineage mutations were obtained from the outbreak.info database^33^ with a threshold of 75%. Lineages that shared the same Spike sequence were merged. The proportion of other lineages that share the same Spike sequence is included within each selected lineage.

### Sequence alignment and phylogenetic reconstruction

The phylogenetic analysis includes the 21 selected lineages and other XBB sub-lineages with a minimum sequence number of 100. The consensus Spike mutations of each lineage were obtained from the outbreak.info database with a threshold of 75%. The amino acid sequence of the Spike protein of each lineage was obtained by replacing each mutation on the reference sequence (WIV04, EPI_ISL_402124). Multiple sequence alignment was performed using MAFFT (v7.453)^34^. Phylogenetic reconstruction was performed with the MEGA X (v10.1) using the minimum evolution method^35^, lineages that shared the identical sequence were merged on the tree. Alignment gaps were pairwise deleted.

### Quantitative and statistical analysis

The statistical analyses for the pseudovirus virus neutralization and antibody binding assessments were performed using GraphPad Prism for calculation of mean value for each data point. Each specimen was tested in triplicate. Antibody neutralization IC_50_ and binding EC_50_ values were calculated using a five-parameter dose-response curve in GraphPad Prism. For comparing the serum neutralization titers and binding titers, statistical analysis was performed using Multiple Mann-Whitney tests. Two-tailed p values are reported. No statistical methods were used to determine whether the data met assumptions of the statistical approach.

## Notes

### Competing Interest Statement

The authors have declared no competing interest.

