## Supplemental Table 1 for "Enhanced neutralization of SARS-CoV-2 XBB sub-lineages and BA.2.86 by a tetravalent COVID-19 vaccine booster"

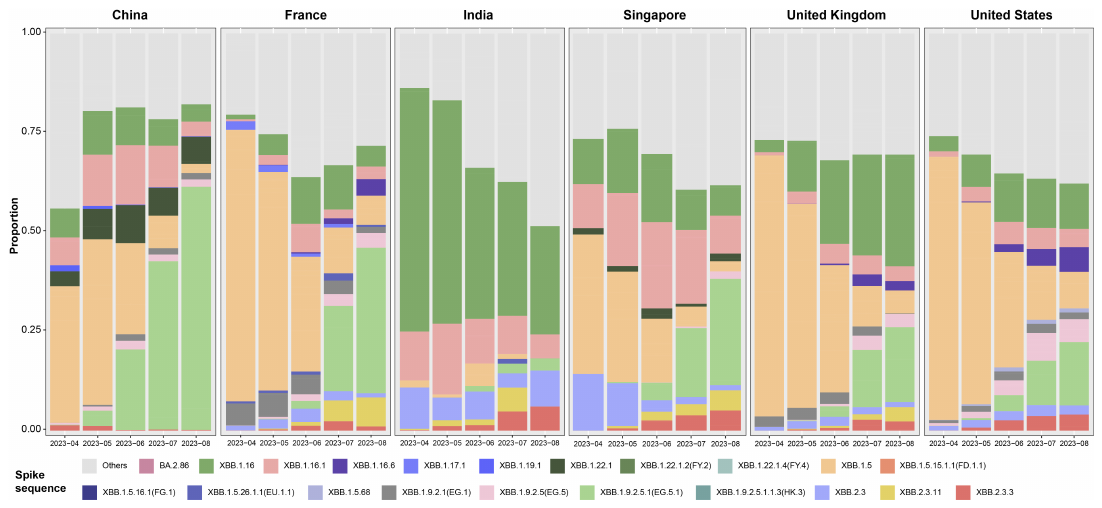
 **Supplementary Figure 1.** Prevalence of Omicron subvariants based on all the sequences available on GISAID from China, France, India, Singapore, the United Kingdom, and the United States, collected between March 15th, 2023, and August 15th, 2023.

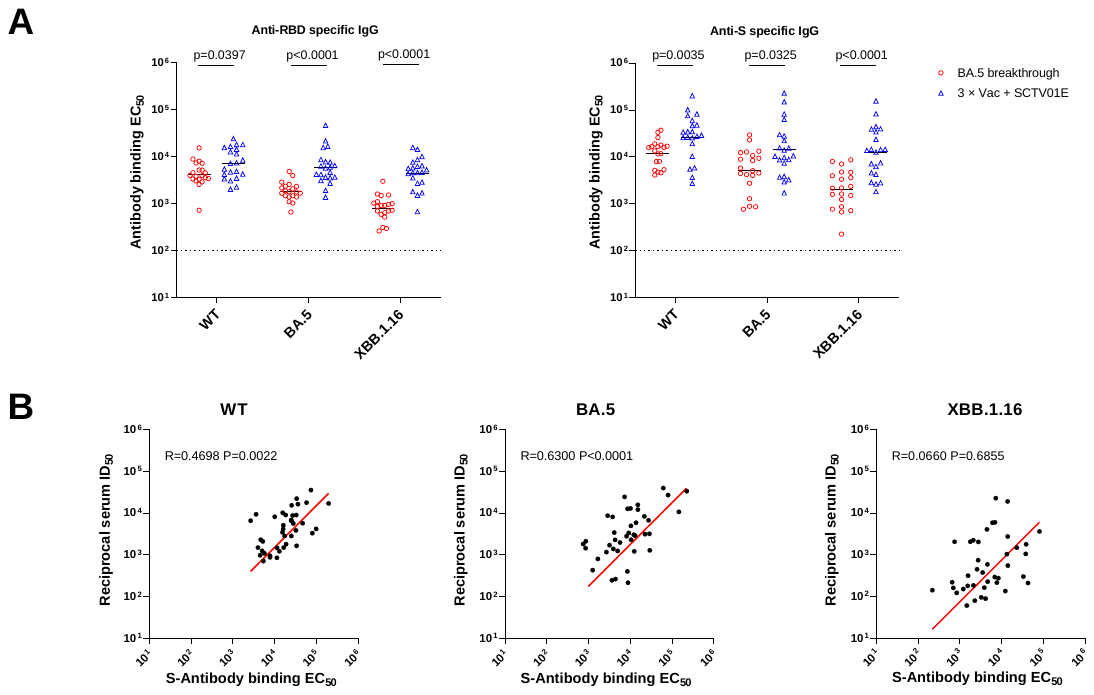

**Supplementary Figure 2. Binding antibody titers against the WT, BA.5, and XBB.1.16 viruses.** (A) Comparative analysis of anti-RBD/S IgG levels between BA.5 breakthrough infection and SCTV01E booster groups. (B) The correlation between anti-S IgG and neutralization ID_50_ combining data from BA.5 breakthrough infection and SCTV01E booster groups. The r value represents the correlation coefficient. Statistics were calculated using Spearman’s rank correlation.

**Supplementary Figure 3.** The full strategies for gating RBD-specific memory B cells (A) and S-specific T cells (B) in flow cytometry.

**Supplementary Table S1. Baseline characteristics of enrolled participants**

|  | 3×CoronaVac individuals(n=11) | 3×Vac+SCTV01E individuals(n=21) | BA.5 breakthrough infection individuals(n=22) | BF.7 breakthrough infection individuals(n=14) | XBB breakthrough infection individuals(n=24) |
| --- | --- | --- | --- | --- | --- |
| Age(years),  median(range) | 40.7(27-50) | 51.0(24-77) | 31.6(22-47) | 40.1(34-50) | 35.3(19-63) |
| Male,n(%) | 6(54.54%) | 16(76.19%) | 13(59.09%) | 4(28.6%) | 13(54.16%) |
| BMI(kg/m^2^),  mean(SD) | 24.8(3.5) | 23.2(3.2) | 23.0(3.9) | 23.9(3.3) | 23.0(3.4) |
| Breakthrough infections months after the last Coronavirus vaccines, median(range) | N/A | N/A | 10.8(2-19) | 10.6(12-14) | 14.5(3-24) |
| Comorbidities(%) | N/A | N/A | N/A | N/A | N/A |
| Any, n(%) | 0(0%) | 0(0%) | 0(0%) | 0(0%) | 0(0%) |
| HTN, n(%) | 0(0%) | 5(23.80%) | 0(0%) | 0(0%) | 2(8.33%) |
| CAD, n(%) | 0(0%) | 0(0%) | 0(0%) | 0(0%) | 0(0%) |
| DM, n(%) | 0(0%) | 2(9.52%) | 0(0%) | 0(0%) | 0(0%) |
| NAFLD, n(%) | 3(27.27%） | 3(14.28%) | 0(0%) | 0(0%) | 1(4.16%) |
| Hyperlipidemia, n(%) | 0(0%) | 1(4.76%) | 0(0%) | 0(0%) | 1(4.16%) |
| Obesity, n(%) | 0(0%) | 1(4.76%) | 2(9.09%) | 0(0%) | 0(0%) |
| Arrhy, n(%) | 0(0%) | 0(0%) | 0(0%) | 0(0%) | 0(0%) |
| Asthma, n(%) | 0(0%) | 0(0%) | 0(0%) | 0(0%) | 0(0%) |
| Rhinitis, n(%) | 0(0%) | 0(0%) | 0(0%) | 0(0%) | 0(0%) |
| Urticaria, n(%) | 0(0%) | 0(0%) | 0(0%) | 0(0%) | 0(0%) |

BMI, body mass index.CAD, coronary artery disease. HTN, hypertension. DM, diabetes mellitus. Arrhy, arrhythmia, NAFLD, non-alcoholic fatty liver.
